# Tracking two decades of global dengue virus lineage dynamics

**DOI:** 10.64898/2026.10.02.756263

**Authors:** Samantha Good, Abigail J. Porzucek, Yi Ting Chew, Rafael Lopes, Naomi Rankin, Christine V. F. Carrington, Eng Eong Ooi, Anderson F. Brito, Verity Hill, Nathan D. Grubaugh

## Abstract

Dengue virus (DENV) is genetically diverse, consisting of four antigenically distinct serotypes and extensive sub-serotype variation that can influence epidemiology and control. As DENV continues to expand globally, there is an urgent need to characterize the dynamics of distinct DENV genetic lineages, including their persistence, establishment, and spread. Using a standardized DENV lineage nomenclature system and 37,828 publicly available DENV genomes sampled from 2000–2024, we classified sequences across nine global regions to construct a comprehensive resource of regional lineage distributions over time. We found marked regional differences in lineage diversity, with higher lineage richness in Southeast and East Asia than in the Americas. We further showed that COVID-19-related disruptions were associated with decreased regional lineage persistence and detection of new lineages during 2020–2021, followed by rebounds after 2022 and, in some regions, substantial lineage turnover. Finally, we identified a small set of globally expansive (“cosmopolitan”) lineages that are over-represented in available sequences, including five first-order minor lineages detected in at least seven of nine regions: 3III_B.3, 2II_F.1, 2II_A.1, 1III_A.3, and 1I_K.3. Together, these results demonstrate how standardized lineage classification enables global tracking of emerging DENV diversity and supports targeted genomic surveillance.

## Introduction

Dengue virus (DENV) poses a major and growing global public health burden, with infections occurring annually across more than 100 countries [1,2]. Global dengue cases have increased thirty-fold over the last two decades, from approximately 500,000 reported cases in 2000 to over 14 million in 2024 [3]. A major factor contributing to the difficulty of controlling dengue is the virus’ extensive genetic diversity: At the broadest level, DENV comprises four antigenically distinct serotypes (DENV-1–4) that frequently co-circulate; while prior infection usually confers long-lasting immunity against symptomatic re-infection with the homotypic DENV, it can enhance the severity of subsequent heterotypic infections [4,5]. This creates a dynamic in which shifts in dominant circulating serotypes can drive dengue outbreaks on roughly 3–8 year cycles [6–9]. Sub-serotype DENV genetic diversity can further alter transmission dynamics and the effectiveness of control measures through antigenic divergence or differences in intrinsic viral fitness [10–20]. Thus, as dengue epidemic waves are frequently driven by the emergence or introduction of distinct viral variants, there is a critical need to characterize within-serotype DENV lineage dynamics at a global scale.

To address this need, we recently developed a phylogenetics-informed DENV lineage nomenclature system (dengue-lineages.org)[21]. This system provides a hierarchical framework for classifying DENV diversity below the serotype level into genotypes, major lineages, and minor lineages, enabling consistent and reproducible comparisons of viral diversity across studies and regions. The system has already proven valuable in tracking the introduction and rapid spread of a new DENV-3 lineage (3III_B.3.2) across the Americas [22–24], characterizing outbreaks in the Caribbean associated with a lineage (2II_F.1.1.5) exhibiting reduced sensitivity to a DENV antiviral [25], and documenting a DENV-1 clade replacement (1I_E.1 by 1I_K.2) in southeastern coastal China during the COVID-19 pandemic [26]. More broadly, characterizing the spatial and temporal patterns of lineage persistence and establishment can reveal how expanding travel networks, urbanization, climate change, and other global disruptions shape DENV genetic diversity worldwide [27]. Applying this framework at a global scale therefore represents a critical next step toward understanding the forces that drive DENV lineage dynamics and improving our capacity to anticipate local outbreak risk, design effective surveillance systems, and inform the deployment of control strategies.

Here, we apply the standardized DENV lineage classification system to 37,828 openly available DENV genomes sampled from 2000 to 2024 to characterize global and regional patterns of lineage diversity, persistence, and spread. We first describe the geographic and temporal distribution of available genomic data and classify sequences by region, year, serotype, and lineage to construct a comprehensive resource for studying DENV epidemiology across nine global regions. We then quantify annual lineage diversity across regions and examine how lineage persistence and new lineage establishment have varied over time. Using these metrics, we investigate how major global and regional disruptions, including the COVID-19 pandemic and the 2015–2016 Zika epidemic, affected DENV lineage dynamics at the regional level. Finally, we identify the lineages with the broadest global spread and highlight those with expanding geographic distributions, providing a framework for ongoing surveillance of emerging DENV lineages worldwide.

## Results

### Global coverage of DENV genomic data

To characterize the global dynamics of DENV lineages, we downloaded all open access DENV sequences from 2000–2024 (**Fig 1**). After data filtering (**Methods**), we categorized 37,828 sequences by geographic region, showing that the data represent broad coverage across the Americas and Asia (North America = 3,088 sequences; South America = 4,950; East Asia = 6,787; Southeast Asia = 16,120; South Asia = 3,710; Pacific = 1,972) but sparser coverage in Africa (936), Europe (210), and the Middle East/North Africa (55; **Fig 1a**). Overall, the total number of available DENV sequences per year increased from 2000–2019, following trends of increasing numbers of reported cases and also likely expanded sequencing capacity (**Fig 1b**). The number of available sequences since 2020, however, have not since reached the pre-COVID-19 pandemic peak, even though the most dengue cases were reported in 2023 and 2024. This may represent a recent de-emphasis in DENV genomic surveillance, open data sharing, or an extended lag in uploading sequences to public repositories. Relative to the number of reported dengue cases, we estimate an overall mean annual sequencing rate of ∼0.08% (sequences/case, 95% CI: 0.07–0.09%), with a higher annual proportion from Asia (East Asia: 4.14% [95% CI: 2.28–6.00%]; Southeast Asia: 0.14% [95% CI: 0.11–0.18%]; South Asia: 0.11% [95% CI: 0.07–0.16%]) than the Americas (North America: 0.06% [95% CI: 0.04–0.08%]; South America: 0.02% [95% CI: 0.01–0.02%]; **Fig 1b**). Despite these discrepancies, we show that the overall DENV serotype proportions are relatively consistent between regions, with DENV-1 and DENV-2 being the most common (**Fig 1c**). Overall, there is sufficient open access DENV genomic data to categorize the sub-serotype lineage dynamics from most regions.

**Fig 1.**
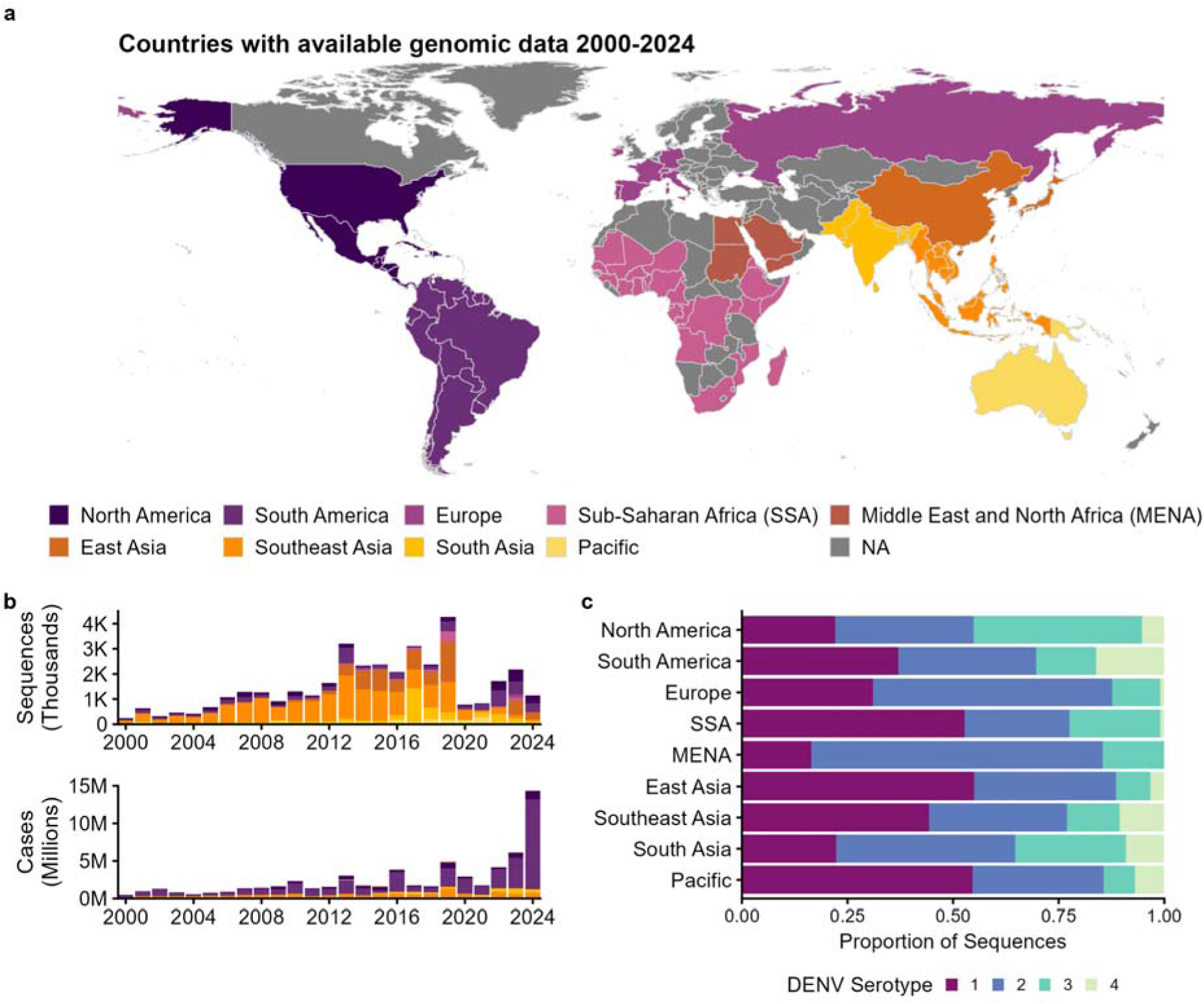
Global distribution of DENV genomic data from 2000-24. (**a**) Global map highlighting the countries and territories reporting DENV sequences from the 37,828 total genomic dataset compiled from 2000-24, colored by region. When possible, DENV sequences from travelers and/or oversea colonies/territories are placed at the origin of infection, not the reporting country. However, there are instances, especially in Europe (e.g. Ireland, Russia) that are likely travel associated but the infection origin is missing. Thus data from Europe should be interpreted with caution. Separate polygons for Hawaii and French Guiana were extracted and re-coded as “Pacific” and “South America” to fix color differences. (**b**) Total annual DENV sequences (top) and reported dengue cases from OpenDengue[28] (bottom) colored by region shown in panel (a). (**c**) The 37,828 DENV sequences separated by region and color b DENV serotype, DENV-1-4. The dataset is available in **S1 Dataset**.

### Major lineage dynamics across geographical regions

Using the new DENV lineage system [21], we classified the 37,828 global sequences by region, year, and to the major lineage level (e.g., serotype 1, genotype 1, major lineage A [1I_A]) when possible to explore the spatial-temporal differences in diversity (**Fig 2, S1 Fig**). These distributions are meant to be a community resource for investigating the epidemiology of DENV lineages, including for studying outbreaks and patterns of spread (**S1 Dataset**).

**Fig 2.**
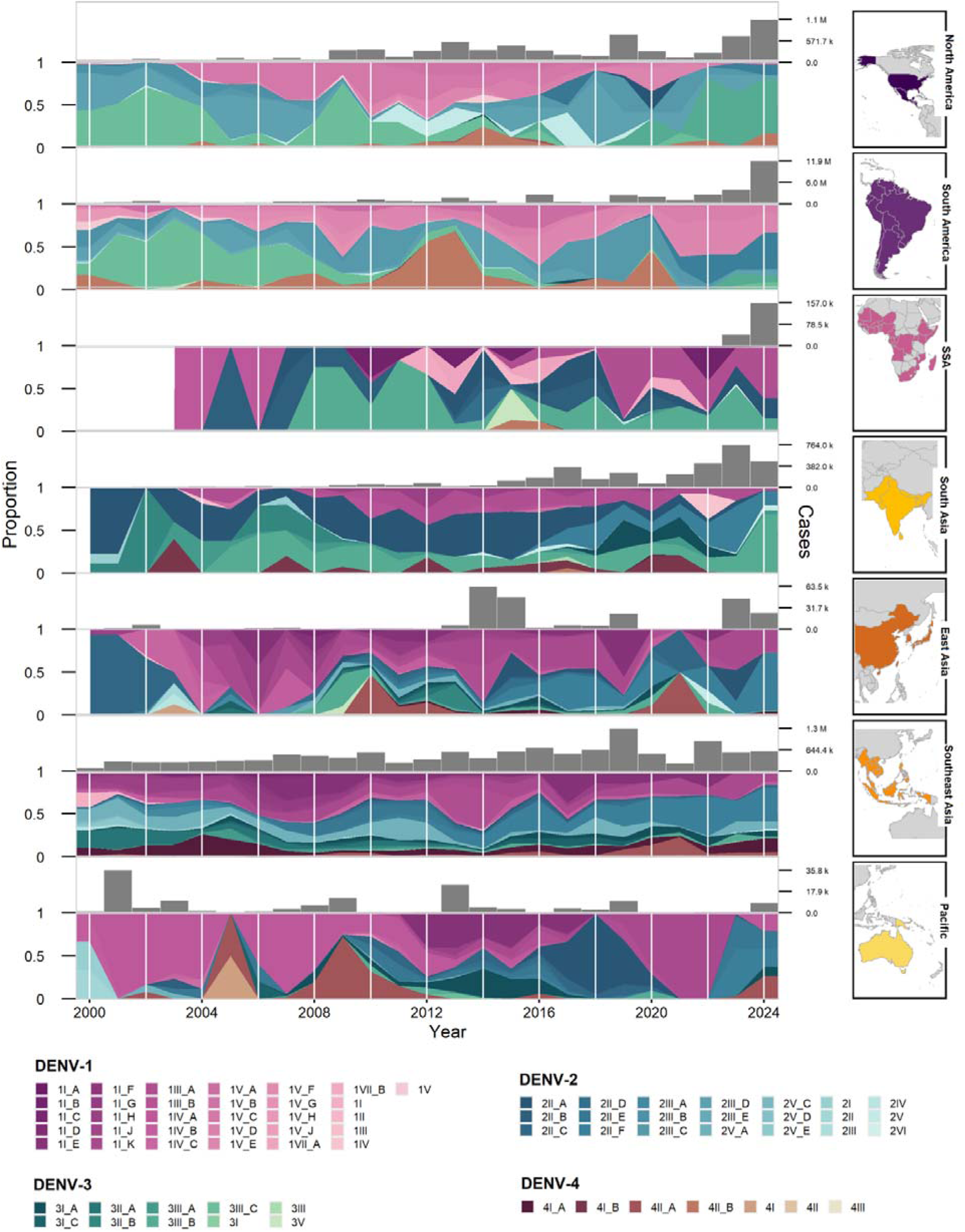
Dynamics of DENV major lineages associated with dengue cases. Stacked area plots showing the proportions of DENV genotypes and major lineages (e.g., 1I_A = DENV serotype 1, genotype 1, major lineage A) for all serotypes, sorted by year from 2000-24 and geographical region (Europe and Middle East/North Africa shown in **S1 Fig**), along with reported yearly dengue cases from each region from OpenDengue[28] shown above in grey. The maps show the countries with available DENV genomic data contributing to the regional proportions. Not all genotypes and major lineages are visible in this plot because their proportions were negligible. The dataset is available in **S1 Dataset**.

To convey how our constructed lineage dynamics help to better understand epidemiological events (**Fig 2**), we provide an example from each region that is supported by published studies. In North America, the re-emergence of DENV-3 since 2022 was associated with the introduction of a new major lineage, 3III_B, into Cuba [29–31], which subsequently spread throughout the Americas where it now competes with the previous dominant DENV-3 lineage, 3III_C [22–24,32–34]. DENV-4 is often detected at low proportions (**Fig 1c**), but the two large peaks of 4II_B in South America capture the 2013 outbreak in Brazil (1.4 million cases reported) [35–37], and the subsequent emergence in Paraguay during 2020 [38,39]. The most dominant lineages in Africa, e.g., 1I_K, 2II_A, and 3III_B, are historically associated with lineages in Asia compared to the Americas, suggesting more opportunities for DENV spread between those regions [40,41]. The dominant circulating DENV-3 lineage in South Asia is 3III_B, but a 2019 outbreak in Sri Lanka was associated with the emergence of 3I_A known to circulate throughout Asia [42]. While DENV-1, specifically lineage 1I_K, continuously circulates in East Asia, a large outbreak in Guangzhou, China in 2014 was associated with the invasion of a new lineage, 1III_A [43–45]. Prior to the outbreak in China, the DENV-1 lineage 1III_A emerged in Southeast Asia to cause an “unprecedented” outbreak of 40,508 confirmed dengue cases in Singapore from 2013-14 [46,47]. After a decade of DENV-1 dominance in the Pacific, DENV-4 lineage 4II_A started spreading across the islands, leading to outbreaks in French Polynesia and New Caledonia in 2009 [48,49]. The rise in autochthonous cases in France in 2023–24 was linked to travel-associated cases of lineage 2II_F linked to the French Caribbean islands (**S1 Fig**) [50]. Dengue is endemic to areas within the Middle East-North Africa region; however, with only 55 sequences from the region, there is limited reliability in interpreting the DENV lineage distributions (**S1 Fig**). Taken together, these examples demonstrate how DENV lineages can be used to investigate outbreaks and patterns of spread.

### Regional differences in annual lineage diversity

Constructing long-term trends in DENV major lineage frequencies (**Fig 2**) also allows us to estimate overall genetic diversity to further explore differences in regional DENV epidemiology (**Fig 3**). We evaluated annual DENV major lineage alpha diversity in two ways: (*1*) richness, the number of distinct major lineages (**Fig 3a-b**), and (*2*) heterogeneity, the interaction between lineage richness and evenness measured by the Gini-Simpson Index (**Fig 3c-d**). We estimated the highest lineage richness in Southeast Asia (mean: 22.2 lineages/year, 95% CI: 19.8–24.7), followed by East Asia, which also exhibited the highest interannual variability (mean: 11.8, CI: 95% 8.5–15.2; **Fig 3a**). Notably, lineage richness in these Asian regions exceeded that of North America (mean: 7.6, CI: 95% 6.7–8.6) and South America (mean: 9.8, 95% CI: 9.1–10.5), despite the latter two regions reporting the highest annual dengue case counts (**Fig 1b**). Temporally, we found no clear trend of increasing lineage richness in any region (**Fig 3b**), despite numerous reports of new lineage introductions. In fact, lineage richness in Southeast and East Asia peaked in 2005 (31 major lineages) and 2013 (25 major lineages), respectively, followed by a marked decline in both regions beginning in 2020. Although we again estimated the highest average Gini-Simpson Index in Southeast Asia, regional and temporal differences in lineage heterogeneity were less pronounced than those observed for richness (**Fig 3c-d**). Overall, DENV lineage diversity was greatest in Southeast Asia, though each region exhibited periods of variable diversity that may reflect underlying DENV transmission dynamics or other epidemiological factors.

**Fig 3.**
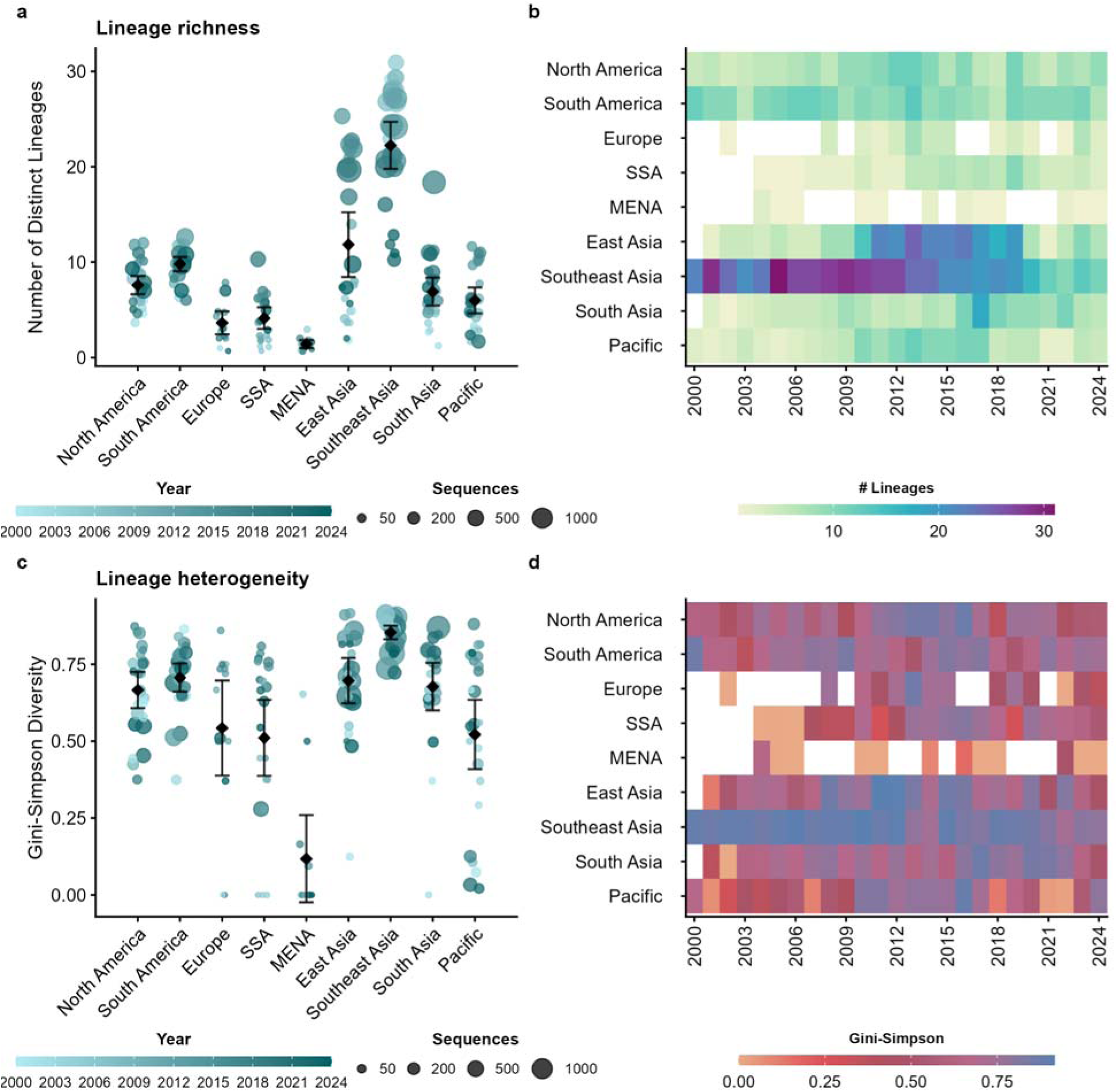
DENV lineage diversity is the highest in Asia. (**a**) Lineage richness represented as the total number of unique major lineages per region and year derived from Fig. 2 from 2000-24, shown in a color gradient, with the circle size representing the number of sequences in the dataset. The mean and 95% CI are shown for each region. (**b**) Same lineage richness data shown in (a) visualized as a heatmap for each region and year with the number of unique lineages shown using a color gradient to better compare the temporal trends. (**c**) Lineage heterogeneity, the interaction between lineage richness and evennes measured by the Gini-Simpson Index per region and year shown in a color gradient, with the circle size representing the number of sequences in the dataset. The mean and 95% CI are shown for each region. (**d**) Same lineage heterogeneity data shown in (c) visualized as a heatmap for each region and year with the number of unique lineages shown using a color gradient to better compare the temporal trends.

We focused our diversity estimates on the major lineages as the minor lineages are more prone to sampling biases. Despite that, biases remain, especially because the overall DENV genome availability varied considerably across regions, with a general decrease in the proportion of sequenced cases after 2019, which impacts the estimates of all lineage classifications (**Fig 1b**). Demonstrating this, we found a positive correlation between the number of sequences per year and the number of distinct major lineages detected in most regions (**S2 Fig**). To address this, we randomly subsampled our data to 50 sequences per year per region and estimated lineage richness (**S2 Fig**). Although this subsampling resulted in more years for which diversity could not be estimated, the overall trends observed in the full dataset were consistent with those in the subsampled data.

### Disruptions to lineage persistence and establishment during the COVID-19 pandemic

The temporal variations we detected in regional DENV lineage diversity (**Fig 3**) may reflect changes in human behavior, from local mobility [51,52] to international travel [27,53,54], that can alter DENV transmission and spread. To investigate these trends, we estimated two metrics for each year from 2010–2024: (*1*) lineage persistence (the percentage of major lineages also detected in the previous year) and (*2*) new lineage establishment (the percentage of total major lineages that were newly detected; **Fig 4**) for each region, excluding the Middle East-North Africa and Europe regions with not enough data. Where we identified gaps in lineage detection between years we assumed lineages remained present throughout, rather than lost and subsequently reintroduced. Across all years and regions, we estimated that 87.9% (95% CI: 83.5–92.3%) of lineages detected in one year persisted into the next, with the highest rates in Southeast Asia (mean: 87.7%, 95% CI: 82.6–92.8%) and South America (mean: 86.2%, 95% CI: 79.0–93.3%) and the lowest in the Pacific (mean: 72.4%, 95% CI: 61.5–83.3%; **Fig 4A**, **S1 Table**). We further estimated that 15.7% (95% CI: 12.6–19.0%) of lineages detected were new to that region and year, with the highest rates of new lineage establishment in Sub-Saharan Africa (mean: 22.1%, 95% CI: 10.8–33.4%) and the Pacific (mean: 21.4%, 95% CI: 8.5–34.4%) and the lowest in Southeast Asia (mean: 6.5%, 95% CI: 3.3–9.8%; **Figure 4B**, **S2 Table**). Although some regional differences may reflect sampling biases, we can nonetheless conclude that low-level lineage emergence is a constant process happening in the background of high rates of lineage persistence.

**Fig 4.**
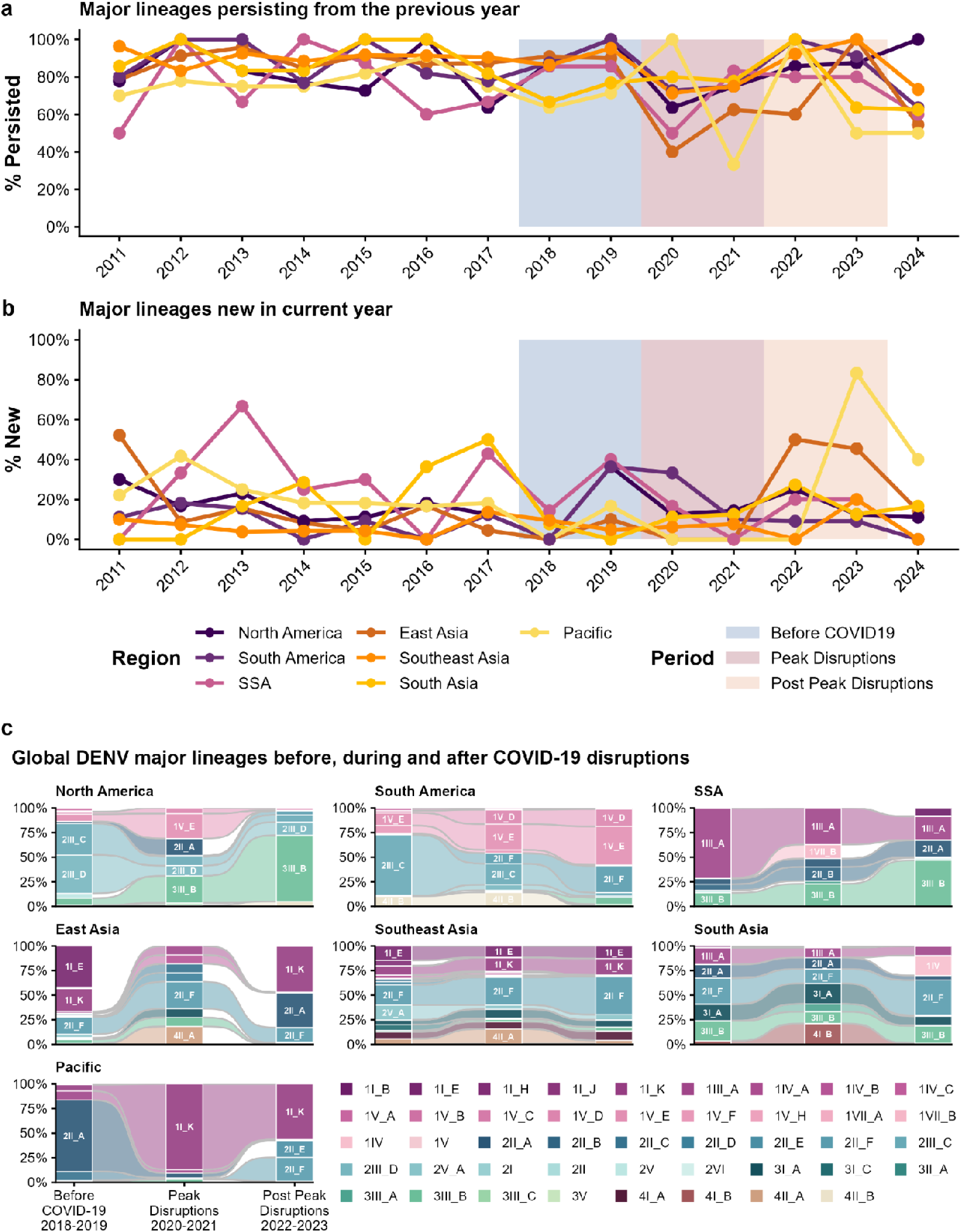
Impacts of COVID-19 disruptions on regional DENV lineage dynamics. (**a**) Line plots showing regional lineage persistence, i.e. the percentage of major lineages also detected in the previous year, colored by region. Two-year periods before the COVID-19 pandemic (2018-19), during the period of peak COVID-19-related disruptions (2020-21), and after the peak disruptions (2022-23) are highlighted for reference. (**b**) Line plots showing new lineage establishment, ie. the percentage of total major lineages that were newly detected, colored by region. Two-year periods around the COVID-19 pandemic are highlighted as described in (a). (**c**) Sankey alluvial plots showing the proportions of DENV genotypes and major lineages per two-year period before (2018-19), during (2020-21), and after (2022-23) the peak disruptions during the COVID-19 pandemic per region. Only the lineages with the top 10 highest proportions are labeled in the plot. The lineage persistence and establishment data can be found in **S1 Table** and **S2 Table**, respectively.

Disruptions during the COVID-19 pandemic, such as reduced time spent in non-residential areas and decreased global air travel, has been linked to local decreases in DENV transmission [55–59] and altered lineage dynamics [26,60,61], perhaps explaining the decline in lineage richness we detected in some regions beginning in 2020 (**Fig 3**). We reasoned that comparing temporal measures lineage persistence and establishment within regions would be less susceptible to systematic error than making between region comparisons, implying that changes during and after the pandemic would likely represent real epidemiological processes. We found that within-region lineage persistence averaged 84.8% (95% CI: 81.8–87.7%) from 2010–2019, declining significantly to 68.5% (95% CI: 58.4–78.7%; *p* = 0.005) during the peak period of COVID-19-related disruptions in 2020–2021, before rebounding to 78.8% (95% CI: 70.3–87.2%) from 2022–2024 as the most restrictive policies were lifted (**Fig 4a**, **S1 Table**). Although we found new lineage establishment to be considerably more variable, ranging from 0–83% across individual years, we nonetheless detected significant differences across the same periods. We estimated that the percentage of newly established lineages averaged 16.0% (95% CI: 12.2–19.8%) from 2010–2019, decreased to 8.9% (95% CI: 3.5–14.2%; *p* = 0.031) during 2020–2021, and increased to 19.8% (95% CI: 10.4–29.2%) from 2022–2024 (**Fig 4b**, **S2 Table**).

These changes in lineage persistence and establishment were not consistent across regions, however, and produced notable differences in DENV lineage frequencies during and after the COVID-19 pandemic (**Fig 4c**). In East Asia, for example, lineage persistence decreased from 88.6% (95% CI: 84.8–92.4%) during 2011–2019 to 40% in 2020 (**Fig 4a**), and when new lineage establishment subsequently exceeded 45% in 2022 and 2023, well above the 2011–2019 mean of 13.3% (95% CI: 1.4–25.3%; **Fig 4b**), the region experienced the most dramatic lineage turnover of any we examined (**Fig 4c**). Specifically, DENV-1I_E was replaced by 1I_K, as previously documented in China [26], and DENV-2II_A rapidly emerged in 2023 to briefly become the dominant serotype 2 lineage in East Asia (**Fig 4c**). In North America, reductions in lineage persistence (2011–2019: 84.6% [95% CI: 74.4–94.8%], 2020: 63.6%, 2021: 75%; **Fig 4a**) and new lineage establishment (2011–2019: 17.5% [95% CI: 8.9–26.0%], 2020: 12.5%, 2021: 14.3%; **Fig 4b**) during the pandemic were less pronounced, resulting in shifts in regional lineage frequencies rather than wholesale replacement, most notably with the emergence of DENV-3III_B.3 (**Fig 4c**). In Southeast Asia, the dominant lineages from before and after the pandemic remained stable (**Fig 4c**), likely a function of the reduction in lineage persistence (2011–2019: 90.7% [95% CI: 87.5–93.8%], 2020: 71.4%, 2021: 75%; **Fig 4a**) mostly impacting low-frequency lineages. Taken together, DENV lineage dynamics were significantly disrupted globally during the COVID-19 pandemic, though the magnitude and nature of these impacts varied by region, likely reflecting differences in the extent and type of pandemic-related disruptions, DENV epidemiology, and genomic surveillance capacity.

Finally, we investigated DENV lineage persistence and establishment in the Americas following the 2015–2016 Zika epidemic, which may have driven a reduction in dengue cases from 2017–2018 and a subsequent large rebound in 2019 [62–64]. In North and South America combined, lineage persistence and new lineage establishment both declined modestly during 2017–2018 (persistence: 2011–16: 87.4% [95% CI: 80.2–94.7%], 2017–18: 79.1% [95% CI: 61.1–97.1%]; establishment: 2011–16: 13.5% [95% CI: 8.0–19.0%], 2017–18: 6.2% [95% CI: 0–17.7%]; **Fig 4a-b, S1-2 Tables**), though neither change was statistically significant.

### Geographical distribution of DENV lineages

Having demonstrated that rates of DENV lineage persistence and establishment vary considerably across regions and time periods (**Fig 4**), suggesting that some lineages are better positioned to spread and establish globally than others, we next investigated which lineages had achieved the greatest global reach (**Fig 5**). We determined the geographical distribution of each major and first-order minor lineage, classified to the first number after the major lineage designation (e.g., 1I_K.3), by counting the number of regions in which each was detected. As expected, major lineages were detected across a greater number of regions (mean: 4.0, 95% CI: 3.3–4.6, n = 52) than minor lineages (mean: 3.0, 95% CI: 2.6–3.5, n = 76; **Fig 5a**; **S3 Table**). We then used these distributions to identify the most geographically distributed (‘cosmopolitan’) lineages, finding that 9 of the 52 major lineages were detected in at least 7 of the 9 regions: 1I_K, 1III_A, 1IV_B, 2II_A, 2II_F, 2III_C, 3I_A, 3III_A, and 3III_B (**S3 Fig**). Although these cosmopolitan lineages represent only 17.3% (9/52) of all classified major lineages, they account for 48.7% (18,433/37,828) of total DENV sequences and encompass 34.2% (26/76) of all designated first-order minor lineages. Specifically, for these cosmopolitan major lineages, their mean cluster size is 2048.1 sequences (95% CI: 882.4–3213.8) and they each give rise to an average of 6.2 minor lineages (95% CI: 2.8–9.7). In comparison, the remaining 43 more geographically restricted major lineages have an average cluster size of 437 sequences (95% CI: 257.8–616.1) and give rise to an average of 4.3 minor lineages (95% CI: 3.1–5.4). These findings indicate that cosmopolitan lineages are not only more geographically widespread but also more genetically diverse and better represented in genomic surveillance data.

**Fig 5.**
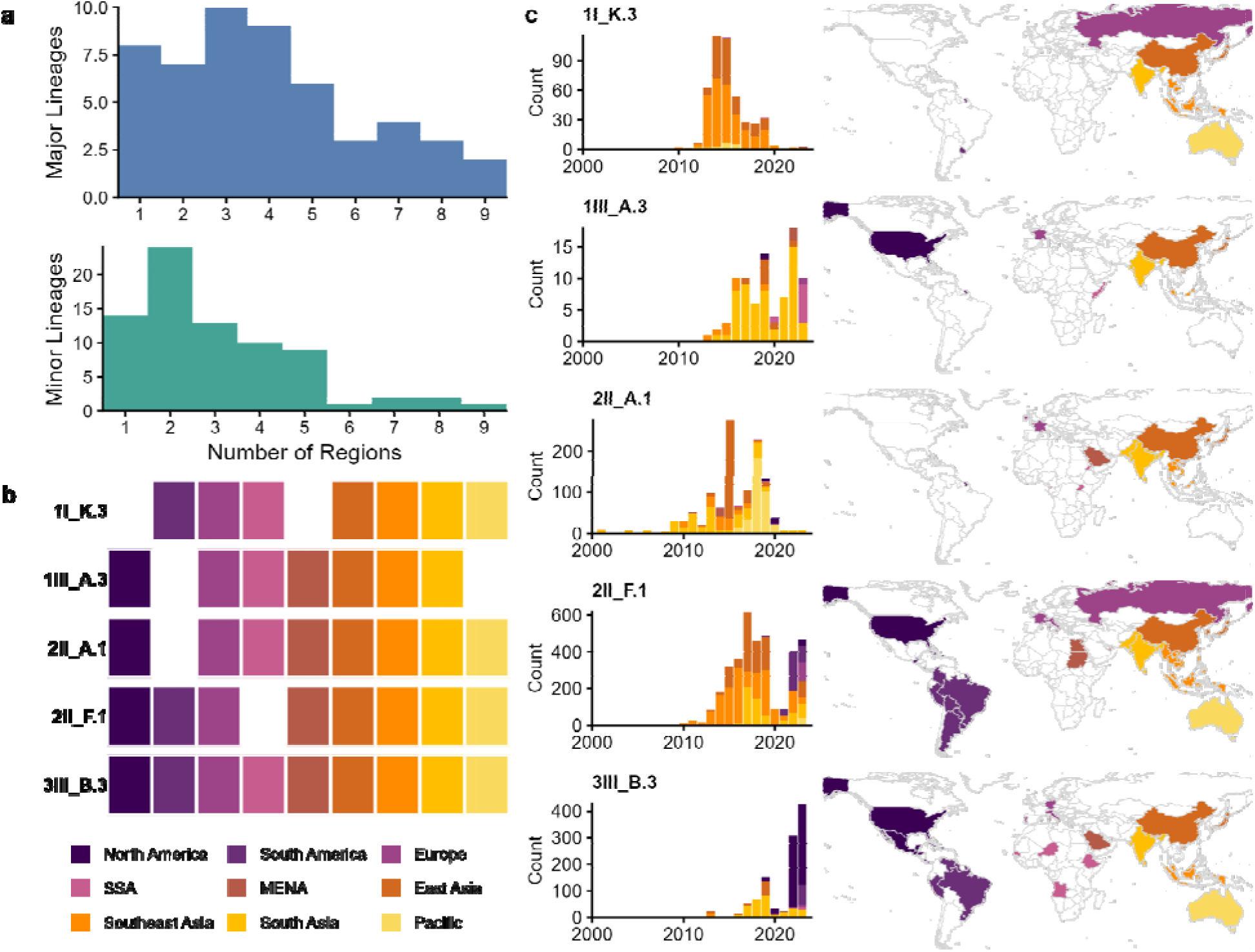
DENV lineages with the greatest global distribution. (**a**) Histogram of the number of geographical regions each major lineage (top) and minor lineage (bottom) has been detected. (**b**) There are five minor lineages detected in 7 or more of the 9 geographical regions, depicted by the colored tiles for each lineage. (**c**) The yearly number of available sequences for each of the five cosmopolitan minor lineages listed in (b), colored by region (left). The maps show the countries, colored by region, that each lineage was detected (right). Note that some of the depicted counties, especially in Europe, likely represent infected travelers instead of local cases.

As some major lineages are quite large and broadly distributed (**S3 Fig**), tracking minor lineages is often more informative for investigating DENV emergence and spread [21]. We found that 5 of the 9 cosmopolitan major lineages also contained first-order minor lineages detected in at least 7 regions: 1I_K.3, 1III_A.3, 2II_A.1, 2II_F.1, and 3III_B.3 (**Fig 5b**). All 5 of these lineages were first detected, and perhaps originated, in Asia before spreading globally (**Fig 5c**). Specifically, 1I_K.3 (452 sequences) was first detected in Southeast Asia (Indonesia) in 2010, subsequently spread throughout Asia and the Pacific with sporadic detections in Sub-Saharan Africa (Tanzania), Europe (Russia, via travel to Thailand [65]), and South America (Uruguay), before declining in global occurrence after 2020 [43,66]. The lineage 1III_A.3 represents a comparatively small cluster (85 sequences) first detected in South Asia (India) in 2002 with frequent detections throughout Asia and sporadic detections in the Middle East-North Africa (Israel), Sub-Saharan Africa (Somalia), and North America (USA, via travel, and French Caribbean Islands) [29,67,68]. The 2II_A.1 lineage (1,192 sequences) pre-dates our dataset, circulates broadly throughout Asia and the Pacific, and has recently been detected across the Middle East, Africa, Europe, and North America, particularly in French-speaking countries [69–71]. We estimate that 2II_F.1 is among the most successful lineages in our dataset (4,058 sequences): first reported from Southeast Asia (Brunei and Singapore) in 2005, it expanded rapidly throughout Asia during the mid-2010s before emerging globally across Africa and the Americas [50,72–75]. Finally, lineage 3III_B.3 (1,414 sequences) first circulated in South and Southeast Asia before rapidly expanding across the Americas, primarily the Caribbean, and spreading to parts of Africa and Europe [29–32,34,76]. Our results thus emphasize the continued importance of DENV genomic surveillance to better track and respond to the global spread of potentially consequential lineages.

## Discussion

Our study provides a global view of DENV lineage dynamics over two decades of publicly available genomes. By analyzing DENV diversity below the serotype level, we reveal epidemiological structure that serotype-only summaries can miss, including regional differences in lineage diversity, the balance of lineage persistence versus newly detected lineages over time, and the subset of cosmopolitan lineages that have spread broadly across regions. Together, our results demonstrate the utility of a standardized, phylogenetics-informed lineage nomenclature for tracking emerging DENV diversity and strengthening genomic surveillance frameworks [21].

A key finding is that DENV lineage diversity is substantially higher in Southeast Asia (and, to a lesser extent, East Asia) than in the Americas, even though the Americas report the highest case burdens [77]. This pattern is consistent with the long-recognized role of Asia as a major reservoir of DENV genetic diversity and sustained hyperendemic transmission, where multiple serotypes and lineages can co-circulate over long periods [78–81]. In our dataset, Southeast Asia also exhibited high lineage persistence and comparatively low proportions of newly detected lineages, consistent with stable, ongoing circulation of a large and diverse lineage pool. In contrast, North and South America showed lower lineage richness but similar or high persistence once lineages were established, reflecting the more recent establishment DENV endemicity punctuated by emerging lineages/serotypes leading to explosive outbreaks [82,83]. Our results underscore the differences in DENV lineage dynamics across regions and provide a baseline to detect future changes that could have significant epidemiological impacts.

We found that the strongest global shifts in temporal lineage diversity occurred during the COVID-19 pandemic period. Other studies have shown that COVID-19-related disruptions, such as the closure of schools, reduced time spent in non-residential areas, and social distancing, led to decreased DENV transmission in some areas [55–59], perhaps questioning the emphasis of residential focused control and providing a mechanism for decreased local lineage persistence and establishment. Further, global disruptions to travel during the pandemic may have reduced lineage introductions, and in some instances, resulted in local lineage turnover when travel resumed [26,60,61,84]. We found evidence that these disruptions impacted DENV lineage dynamics at region scales, specifically a decline in year-to-year lineage persistence and the fraction of lineages newly detected during 2020–2021 which then partially rebounded after 2022. Importantly, the magnitude and character of these shifts varied by region. East Asia displayed especially pronounced turnover, with sharp decreases in persistence followed by elevated proportions of newly detected lineages, consistent with substantial reshaping of lineage composition during and after pandemic disruptions (e.g. [26]). Other regions showed more modest changes, suggesting that the effects of COVID-19-related disruptions on DENV lineage dynamics depend on baseline transmission patterns, connectivity, human behavior (including local policies and culture), and surveillance intensity. We also observed modest declines in persistence and newly detected lineages in the Americas following the 2015–2016 Zika epidemic period, aligning with broader regional decline in reported dengue cases [62,85], meaning that DENV lineages appeared to cryptically persist until dengue rebounded in 2019 [63]. However, these effects were smaller than those observed during the COVID-19 period.

We identified the DENV lineages with the broadest global spread as a way to track “successful” DENV lineages that may have broad fitness advantages. We found an over-representation of globally expansive major lineages in available sequences, and we detected five first-order minor lineages in at least seven of nine global regions. These “cosmopolitan” lineages provide a practical shortlist for ongoing surveillance because they represent repeatedly successful invasions across diverse epidemiological contexts. These include DENV-2II_F.1 and 3III_B.3 that have both recently emerged in the Americas, sometimes being associated with large outbreaks [29,30,34,72,73]. Notably, each of the most geographically widespread minor lineages was first detected in Asian regions before later appearing elsewhere, reinforcing Asia’s central role in shaping global DENV lineage diversity [27].

Several limitations should be considered when interpreting these results. First, the availability of open-access DENV genomes remains sparse relative to case numbers (∼0.08%) and varies substantially by region and year, which can bias lineage detection and estimating accurate proportions. Second, sequencing and data submission are uneven across countries within regions, meaning that regional summaries may be disproportionately shaped by a subset of well-sampled locations (most data come from Brazil, China, Thailand, Vietnam, and Singapore). Because of these issues, we emphasize that temporal comparisons within regions (e.g., persistence and newly detected lineages over time) are generally more robust than absolute between-region comparisons, and we note that many patterns identified here align with published country-level studies. Future work could improve inference by explicitly accounting for surveillance effort (e.g., weighting by case burden and sequencing intensity), expanding access to genomes from under-sampled regions, and further refining geographic attribution where territories and sovereignties and travel-acquired infections complicate regional assignment.

Overall, our study shows that DENV lineage diversity and turnover vary widely across the world and are sensitive to major societal disruptions, while a small subset of lineages achieves broad geographic spread. Tracking globally expanding lineages and linking them to disease outcomes, antigenic divergence, and transmission phenotypes through clinical and experimental studies will strengthen surveillance systems and preparedness for future dengue outbreaks [16].

## Materials and methods

### Dataset construction

Dengue virus sequences by serotype were downloaded on 2025-06-06 from NCBI Virus. Note that we did not use any data from GISAID as we focus on fully open data, and they only started openly hosting DENV sequences in 2023 [86]. We first selected all DENV sequences with sampling dates between 2000 to 2024 (n=41,888). We then excluded sequences with less than 5% genome coverage (505 base pairs), as that is the minimum sequence length required to accurately assign lineages [21], any lab passaged or vaccine strain sequences, and sequences with missing dates or locations.

We assigned DENV lineage assignments using the new nomenclature system (dengue-lineages.org)[21] by uploading the nucleotide FASTA files to NextClade v3.18.1 [87] for clade assignment for each serotype (DENV-1 = 16,298 sequences; DENV-2 = 12,965; DENV-3 = 6,015; DENV-4 = 3,486). Disease of interest and serotype references were autoselected by NextClade. Sequences that successfully received lineage assignments via NextClade were combined with their metadata using Python 3.12.1. The final dataset for analysis was: DENV-1 = 16,088 sequences; DENV-2 = 12,908, DENV-3 = 5,987: DENV-4 = 3,476.

### Dataset Preprocessing

After initial download, we retained the variables: “Accession”, “Isolate”, “Submitters”, “Collection_Date”, “Geo_Location”, and “Country”. NextClade-assigned lineages were joined to the sequence dataset by “Accession”. In data preprocessing new variables, we generated “Year” and “Region” using the extracted year from “Collection_Date” and the “Country” values. States and territories that fall geographically in a different region than their country of sovereignty were reclassified, e.g. “Hawaii” was reclassified as “Pacific” instead of falling under “North America” with the rest of the United States, and “French Guiana” was included with “South America” instead of “Europe”. Additionally each unique variable was checked for consistent spelling. Regions were decided using dengue risk status as well as ecological and epidemiological uniqueness. Chosen regions for this study are: “North America” (including Central America and the Caribbean), “South America”, “Europe”, “Middle East and North Africa”, “Sub-Saharan Africa”, “South Asia”, “East Asia”, “Southeast Asia”, and “Pacific”. When possible, we assigned travel-acquired cases to the location of their travel instead of the country of detection.

Each analysis was performed with minor and major lineage levels. For major lineage classifications we dropped minor lineage nomenclature at the first decimal point. For minor lineages we dropped lineages that did not contain a decimal then collapsed the lineages to their first order decimal, minimizing excess sublineages of the minor lineages.

We acquired annual country-level dengue cases from OpenDengue[28] on 15th November 2025 and then combined case numbers to the region classifications described above.

### Data Visualization

Genomic data analysis was performed in RStudio v4.4.2. We used ggplot2 [88] for all visualizations. For **Fig 2** area plots we used proportions of each lineage for each region and year combination (i.e. South-America, 2000), and we excluded any rows with no proportion data before plotting. To display both case and sequence data in **Fig 2**, we mathematically scaled the cases and added 1 to manually sit case bars on top of the area plots. **S2 Fig** scatterplot fits a straight line through the points using method = “lm”. Geom_smooth was set to display a 95% confidence interval around this line and a Pearson correlation using stat_cor from ggpubr [89].

### Lineage Diversity Analyses

**Fig 3** used a consistent seed set at 42 so the position jitter function within geom_point() would stagger the points the same way each time. **S2 Fig** also uses a seed at 42 to make the subsampling reproducible. We calculated error bars for the richness/heterogeneity plots from each region’s mean number of lineages, the t critical value (df = n-1), multiplied by the standard error. Gini-simpson diversity was calculated on a matrix of lineage proportions in each region-year combination using the vegan package in R [90] with the index set to “simpson”. For subsampled richness analysis, the dataset was grouped by region and year and removed any of these rows where the number of sequences was less than 50. Then a random sample of 50 sequences from region-years with enough sequences to contribute was pulled for the plot without replacement.

### Lineage Persistence and Establishment Analyses

We first imputed lineages for years where that lineage existed the year before and the year after, making the assumption that the lineage must not have been captured in sequencing opposed to disappearing and being reintroduced. Then we calculated lineage persistence by totaling the lineages from the previous year that persisted into the current year and dividing that number by total lineages in the current year. New lineage establishment in the current year was calculated as the lineages not seen in the previous year divided by the total lineages in the current year. **Fig 4** Sankey plots used ggalluvial [91] to calculate the ribbon across year categories.

### Software Packages and Data Availability

Data manipulation used dplyr, stringr and forcats from tidyverse [92]. Shapefiles for maps were downloaded from rnaturalearth [93] and rnaturalearthdata [94]. Additional packages used were cowplot [95], ggh4x [96], grid [97], ltc [98], paletteer [99], patchwork [100], rio [101], scales [102], and sf [103,104]. The genomic data, metadata, lineage proportions, and R and Python scripts can be found at: https://github.com/grubaughlab/2026_paper_DENV-global-lineages

## Acknowledgements

We thank all of the DENV sequencers and data submitters from around that world that make work like this possible. We also acknowledge technical support from D. Simmonds, S. Taylor, P. Jack, P. Carlton, and V. Parsons. This publication was made possible by the National Institute of Allergy and Infectious Diseases of the National Institutes of Health (NIH) under Award Number DP2AI176740 (NDG), the Centers for Disease Control and Prevention of the U.S. Department of Health and Human Services (HHS) Award Number NU50CD300866 (SG), and the Wellcome Trust Award Number 327522/Z/25/Z (VH). The findings and conclusions in this report are those of the authors and do not necessarily represent the official position of, nor an endorsement by, the NIH, CDC/HHS, or the U.S. Government.

## Supporting information

**S1 Dataset** - Figure 2 all lineage proportions (csv)

**S1 Table** - Lineage persistence (word doc)

**S2 Table** - New lineage establishment (word doc)

**S3 Table** - First detection of globally spread lineages (word doc)

**S1 Fig.**
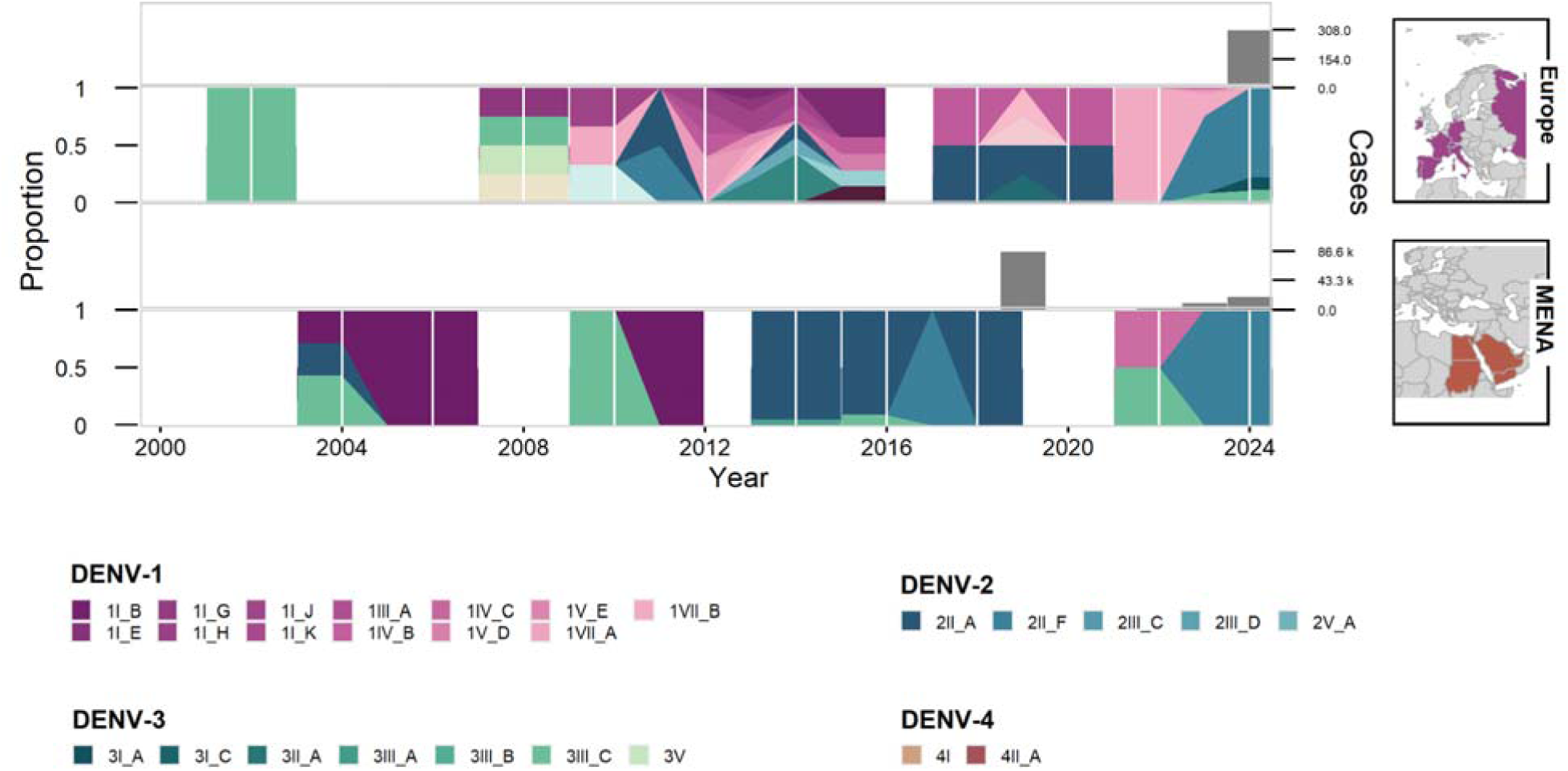
Dynamics of DENV major lineages from Europe and Middle East/North Africa. Stacked area plots showing the proportions of DENV genotypes and major lineages (e.g., 1I_A = DENV serotype 1, genotype 1, major lineage A) for all serotypes, sorted by year from 2000-24 and geographical region Europe or Middle East/North Africa (MENA), along with reported yearly dengue cases from each region from OpenDengue[28] shown above in grey. The maps show the countries with available DENV genomic data contributing to the regional proportions. Not all genotypes and major lineages are visible in this plot because their proportions were negligible. Related to Fig. 2. The dataset is available in **S1 Dataset**.

**S2 Fig.**
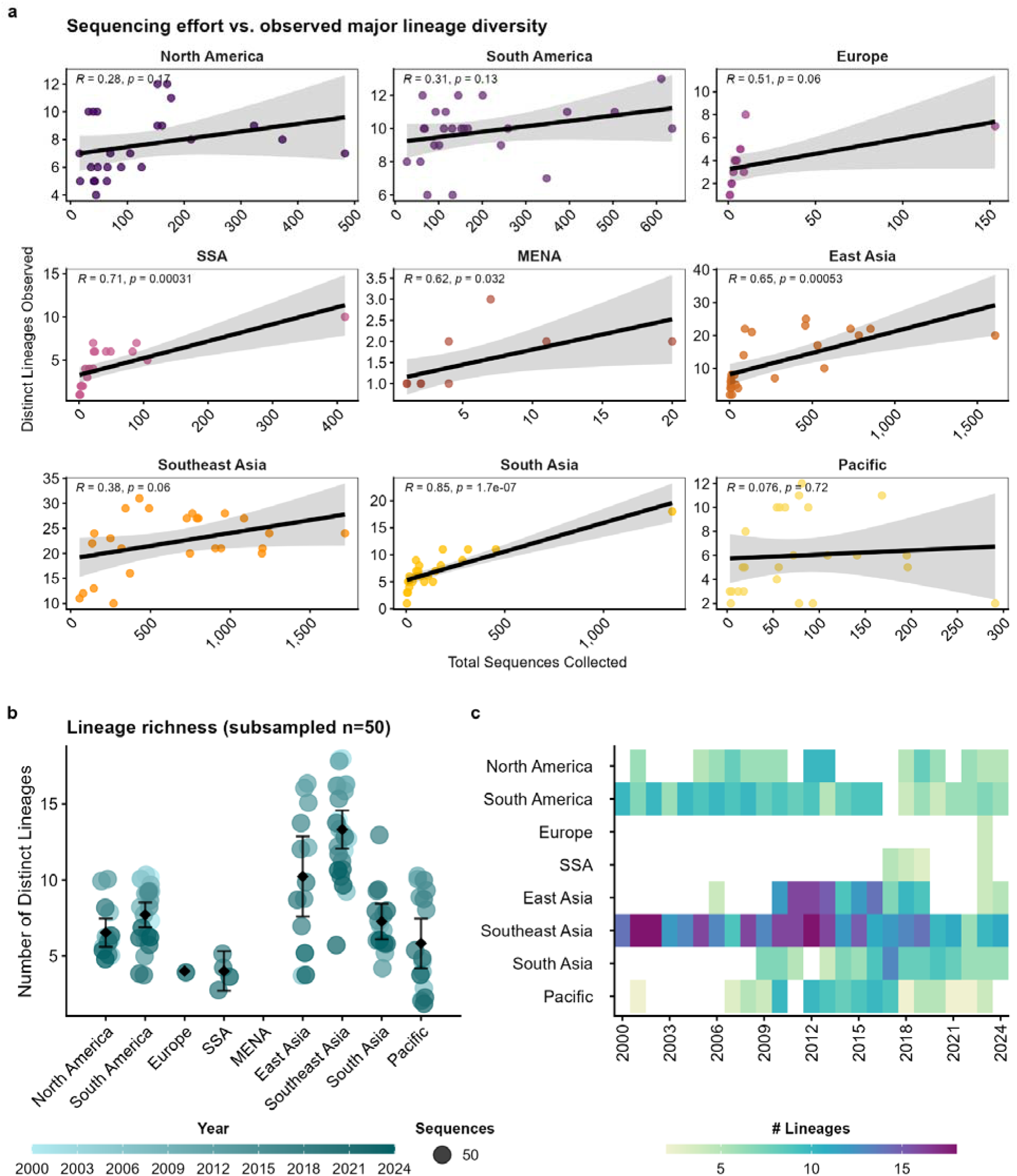
Effects of sampling size on lineage richness estimates. (**a**) Regional Pearson correlation between the number of sequences and distinct lineages detected per year and the 95% CI shown in grey. (**b**) Lineage richness estimates after randomly subsampling 50 sequences per region and year from Fig. 3a, shown in a color gradient representing the year. The mean and 95% CI are shown for each region. (**c**) Same lineage richness data shown in (b) visualized as a heatmap for each region and year with the number of unique lineages shown using a color gradient to better compare the temporal trends. Related to Fig. 3.

**S3 Fig.**
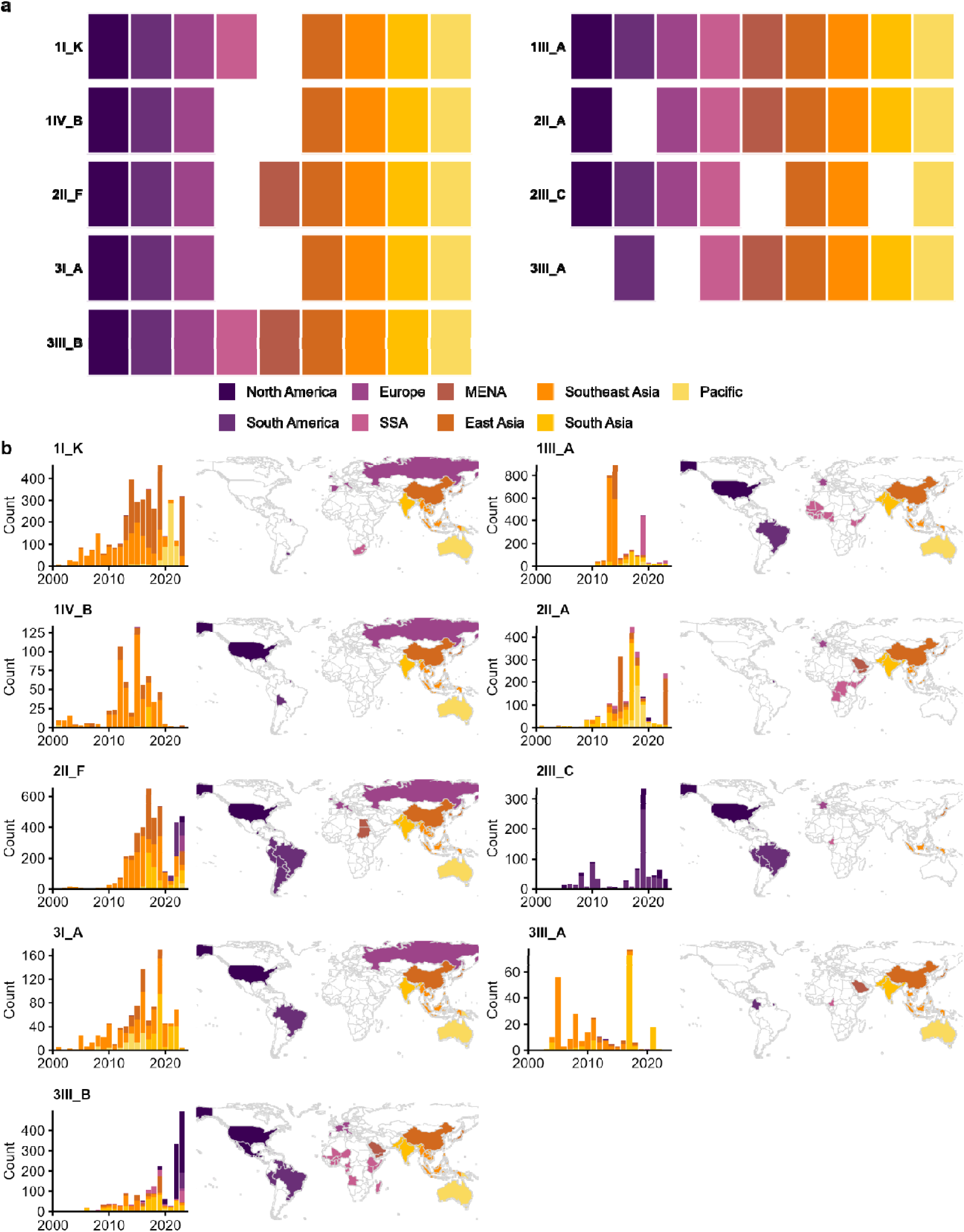
DENV major lineages with the greatest global distribution. (**a**) There are nine major lineages detected in 7 or more of the 9 geographical regions, depicted by the colored tiles for each lineage. (**b**) The yearly number of available sequences for each of the nine major lineages listed in (a), colored by region (left). The maps show the countries, colored by region, that each lineage was detected (right). Note that some of the depicted counties, especially in Europe, likely represent infected travelers instead of local cases. Related to Fig. 5.

## Notes

https://github.com/grubaughlab/2026_paper_DENV-global-lineages

### Competing Interest Statement

The authors have declared no competing interest.

