## Supplementary material for "Tracking two decades of global dengue virus lineage dynamics": S1 Table

S1 Table. Lineage persistence by region and time period

| **2010-2019** | | | | **2020-2021** | | | **2022-2024** | | |
| --- | --- | --- | --- | --- | --- | --- | --- | --- | --- |
| **Region** | **Total Unique Lineages** | **Mean % (95% CI)** | **Median % (IQR)** | **Total Unique Lineages** | **Mean % (95% CI)** | **Median % (IQR)** | **Total Unique Lineages** | **Mean % (95% CI)** | **Median % (IQR)** |
| ***Observed — % Lineages Persisted*** | | | |  | | |  | | |
| North America | 22 | 71.7% (95% CI:  57.4–86%) | 75% (IQR: 50–  85.7%) | 9 | 48.7% (95% CI:  37.3–60.1%) | 48.7% (IQR:  45.8–51.6%) | 10 | 85% (95% CI: 70–  100%) | 80% (IQR: 77.5–  90%) |
| South America | 18 | 84.4% (95% CI:  77.4–91.5%) | 80% (IQR: 77.8–  88.9%) | 13 | 62.2% (95% CI:  47.1–77.4%) | 62.2% (IQR:  58.4–66.1%) | 12 | 81.2% (95% CI: 64–  98.4%) | 90% (IQR: 76.8–  90%) |
| Europe | 19 | 18.8% (95% CI:  1.1–36.4%) | 12.5% (IQR: 0–  34.4%) | 2 | 25% (95% CI: NA–NA%) | 25% (IQR: 25–  25%) | 8 | 64.3% (95% CI: -  5.7–134.3%) | 64.3% (IQR:  46.5–82.2%) |
| MENA | 5 | 0% (95% CI: 0–0%) | 0% (IQR: 0–0%) | - | - | - | 3 | 50% (95% CI: -48–  148%) | 50% (IQR: 25–  75%) |
| SSA | 16 | 57.1% (95% CI:  46.8–67.5%) | 50% (IQR: 50–  66.7%) | 6 | 58.4% (95% CI:  42–74.7%) | 58.4% (IQR:  54.2–62.5%) | 6 | 61.7% (95% CI:  47.4–75.9%) | 60% (IQR: 55–  67.5%) |
| East Asia | 38 | 81% (95% CI:  76.8–85.3%) | 81.8% (IQR: 76–  86.4%) | 8 | 24.6% (95% CI:  4.4–44.9%) | 24.6% (IQR:  19.5–29.8%) | 12 | 44.4% (95% CI: -3–  91.9%) | 50% (IQR: 25–  66.7%) |
| Southeast Asia | 37 | 85.6% (95% CI:  83.3–87.8%) | 85.2% (IQR:  83.3–87.5%) | 17 | 69.4% (95% CI:  68.2–70.6%) | 69.4% (IQR:  69.1–69.7%) | 15 | 74.7% (95% CI:  68.6–80.8%) | 75% (IQR: 72.1–  77.5%) |
| South Asia | 21 | 78.3% (95% CI:  68.2–88.4%) | 80% (IQR: 66.7–  85.7%) | 10 | 77.8% (95% CI:  77.8–77.8%) | 77.8% (IQR:  77.8–77.8%) | 13 | 70.5% (95% CI:  41.6–99.4%) | 57.1% (IQR:  55.8–78.6%) |
| Pacific | 21 | 63.9% (95% CI:  56.6–71.1%) | 66.7% (IQR: 60–  70%) | 6 | 66.7% (95% CI:  1.3–132%) | 66.7% (IQR: 50–  83.3%) | 9 | 66.7% (95% CI: 34–  99.3%) | 50% (IQR: 50–  75%) |
| ***Assumed (gap-filled) — % Lineages Persisted*** | | | |  | | |  | | |
| North America | 22 | 84.6% (95% CI:  76–93.3%) | 83.3% (IQR:  76.9–100%) | 9 | 69.3% (95% CI:  58.1–80.5%) | 69.3% (IQR:  66.4–72.2%) | 10 | 91.1% (95% CI:  82.3–99.9%) | 87.5% (IQR:  86.6–93.8%) |
| South America | 18 | 89.3% (95% CI:  82.4–96.2%) | 87.5% (IQR: 80–  100%) | 13 | 73.8% (95% CI:  71.6–76.1%) | 73.8% (IQR:  73.3–74.4%) | 12 | 84.8% (95% CI:  63.4–106.3%) | 90.9% (IQR:  77.2–95.4%) |

| **2010-2019** | | | | **2020-2021** | | | **2022-2024** | | |
| --- | --- | --- | --- | --- | --- | --- | --- | --- | --- |
| **Region** | **Total Unique Lineages** | **Mean % (95% CI)** | **Median % (IQR)** | **Total Unique Lineages** | **Mean % (95% CI)** | **Median % (IQR)** | **Total Unique Lineages** | **Mean % (95% CI)** | **Median % (IQR)** |
| Europe | 19 | 48.1% (95% CI:  25–71.3%) | 41.6% (IQR:  33.3–54.2%) | 2 | 40% (95% CI: NA–NA%) | 40% (IQR: 40–  40%) | 8 | 64.3% (95% CI: -  5.7–134.3%) | 64.3% (IQR:  46.5–82.2%) |
| MENA | 5 | 50% (95% CI: 19–  81%) | 50% (IQR: 50–  50%) | - | - | - | 3 | 50% (95% CI: -48–  148%) | 50% (IQR: 25–  75%) |
| SSA | 16 | 78% (95% CI:  66.4–89.7%) | 85.7% (IQR:  66.7–87.5%) | 6 | 66.7% (95% CI:  34–99.3%) | 66.7% (IQR:  58.3–75%) | 6 | 73.3% (95% CI:  60.3–86.4%) | 80% (IQR: 70–  80%) |
| East Asia | 38 | 88.6% (95% CI:  85.4–91.8%) | 90% (IQR: 87–  91.3%) | 8 | 51.2% (95% CI:  29.2–73.3%) | 51.2% (IQR:  45.6–56.9%) | 12 | 71.5% (95% CI:  43.4–99.6%) | 60% (IQR: 57.2–  80%) |
| Southeast Asia | 37 | 90.7% (95% CI:  88–93.3%) | 91.3% (IQR:  88.5–92.6%) | 17 | 73.2% (95% CI:  69.7–76.7%) | 73.2% (IQR:  72.3–74.1%) | 15 | 88.5% (95% CI: 73–  104.1%) | 92.3% (IQR:  82.8–96.2%) |
| South Asia | 21 | 86.4% (95% CI:  78.8–94%) | 83.3% (IQR:  81.8–100%) | 10 | 78.9% (95% CI:  76.7–81.1%) | 78.9% (IQR:  78.3–79.4%) | 13 | 75.4% (95% CI:  51.2–99.5%) | 63.6% (IQR: 63–  81.8%) |
| Pacific | 21 | 75.6% (95% CI:  70.6–80.6%) | 75% (IQR: 71.4–  77.8%) | 6 | 66.7% (95% CI:  1.3–132%) | 66.7% (IQR: 50–  83.3%) | 9 | 66.7% (95% CI: 34–  99.3%) | 50% (IQR: 50–  75%) |
