## Supplementary material for "Tracking two decades of global dengue virus lineage dynamics": S2 Table

S2 Table. Newly established lineages by region and time period

| **2010-2019** | | | | **2020-2021** | | | **2022-2024** | | |
| --- | --- | --- | --- | --- | --- | --- | --- | --- | --- |
| **Region** | **Total Unique Lineages** | **Mean % (95% CI)** | **Median % (IQR)** | **Total Unique Lineages** | **Mean % (95% CI)** | **Median % (IQR)** | **Total Unique Lineages** | **Mean % (95% CI)** | **Median % (IQR)** |
| ***Observed — % New Lineages*** | | | |  | | |  | | |
| North America | 22 | 32% (95% CI: 24.4–  39.6%) | 30% (IQR: 25–  40%) | 9 | 27.1% (95% CI: 2–  52.3%) | 27.1% (IQR:  20.7–33.6%) | 10 | 28.8% (95% CI:  7.6–50.1%) | 22.2% (IQR:  18.2–36.1%) |
| South America | 18 | 15.9% (95% CI: 6.9–  24.9%) | 12.5% (IQR:  11.1–20%) | 13 | 35% (95% CI:  25.2–44.8%) | 35% (IQR: 32.5–  37.5%) | 12 | 9.4% (95% CI: -0.9–  19.7%) | 10% (IQR: 5–  14.1%) |
| Europe | 19 | 83.8% (95% CI:  64.8–102.7%) | 93.8% (IQR:  78.1–100%) | 2 | 50% (95% CI: NA–NA%) | 50% (IQR: 50–  50%) | 8 | 59.5% (95% CI:  8.1–110.9%) | 59.5% (IQR:  46.4–72.6%) |
| MENA | 5 | 100% (95% CI: 100–  100%) | 100% (IQR: 100–  100%) | - | - | - | 3 | 50% (95% CI: -48–  148%) | 50% (IQR: 25–  75%) |
| SSA | 16 | 44.2% (95% CI:  27.5–60.8%) | 50% (IQR: 33.3–  50%) | 6 | 8.3% (95% CI: -8–  24.7%) | 8.3% (IQR: 4.2–  12.5%) | 6 | 30% (95% CI: 0.1–  59.9%) | 40% (IQR: 20–  45%) |
| East Asia | 38 | 20.9% (95% CI: 12–  29.9%) | 14.3% (IQR:  13.6–24%) | 8 | 25% (95% CI: -24–  74%) | 25% (IQR: 12.5–  37.5%) | 12 | 59.5% (95% CI:  18.1–101%) | 50% (IQR: 39.3–  75%) |
| Southeast Asia | 37 | 11.1% (95% CI: 7.3–  14.8%) | 11.1% (IQR: 5–  15%) | 17 | 10.4% (95% CI:  6.3–14.5%) | 10.4% (IQR: 9.4–  11.4%) | 15 | 22.2% (95% CI:  5.6–38.8%) | 18.2% (IQR:  14.1–28.4%) |
| South Asia | 21 | 22.9% (95% CI:  10.4–35.4%) | 20% (IQR: 9.1–  42.9%) | 10 | 17.4% (95% CI:  7.8–26.9%) | 17.4% (IQR:  14.9–19.8%) | 13 | 25% (95% CI: 14–  36%) | 27.3% (IQR:  20.8–30.3%) |
| Pacific | 21 | 31.3% (95% CI:  19.8–42.7%) | 27.3% (IQR: 25–  41.7%) | 6 | 0% (95% CI: 0–0%) | 0% (IQR: 0–0%) | 9 | 41.1% (95% CI: -6–  88.2%) | 40% (IQR: 20–  61.6%) |
| ***Assumed (gap-filled) — % New Lineages*** | | | |  | | |  | | |
| North America | 22 | 17.5% (95% CI:  10.2–24.7%) | 16.7% (IQR:  11.1–23.1%) | 9 | 13.4% (95% CI:  11.6–15.2%) | 13.4% (IQR:  12.9–13.9%) | 10 | 16.2% (95% CI:  7.5–24.9%) | 12.5% (IQR:  11.8–18.8%) |
| South America | 18 | 11.4% (95% CI: 3.8–  19%) | 11.1% (IQR: 0–  15.4%) | 13 | 21.6% (95% CI: -  1.2–44.5%) | 21.6% (IQR:  15.8–27.5%) | 12 | 6.1% (95% CI: 0.1–  12%) | 9.1% (IQR: 4.6–  9.1%) |

| **2010-2019** | | | | **2020-2021** | | | **2022-2024** | | |
| --- | --- | --- | --- | --- | --- | --- | --- | --- | --- |
| **Region** | **Total Unique Lineages** | **Mean % (95% CI)** | **Median % (IQR)** | **Total Unique Lineages** | **Mean % (95% CI)** | **Median % (IQR)** | **Total Unique Lineages** | **Mean % (95% CI)** | **Median % (IQR)** |
| Europe | 19 | 62.7% (95% CI:  43.7–81.7%) | 70.8% (IQR:  61.7–77.1%) | 2 | 0% (95% CI: NA–NA%) | 0% (IQR: 0–0%) | 8 | 59.5% (95% CI:  8.1–110.9%) | 59.5% (IQR:  46.4–72.6%) |
| MENA | 5 | 40% (95% CI: 3.3–  76.7%) | 50% (IQR: 0–  50%) | - | - | - | 3 | 50% (95% CI: -48–  148%) | 50% (IQR: 25–  75%) |
| SSA | 16 | 28% (95% CI: 14–  42%) | 30% (IQR: 14.3–  40%) | 6 | 8.3% (95% CI: -8–  24.7%) | 8.3% (IQR: 4.2–  12.5%) | 6 | 13.3% (95% CI:  0.3–26.4%) | 20% (IQR: 10–  20%) |
| East Asia | 38 | 13.3% (95% CI: 3.2–  23.5%) | 8.7% (IQR: 4.5–  15.4%) | 8 | 0% (95% CI: 0–0%) | 0% (IQR: 0–0%) | 12 | 36.6% (95% CI:  14.6–58.6%) | 45.5% (IQR:  29.9–47.8%) |
| Southeast Asia | 37 | 6.4% (95% CI: 3.7–  9.1%) | 4.8% (IQR: 4.2–  9.5%) | 17 | 7% (95% CI: 5.5–  8.4%) | 7% (IQR: 6.6–  7.3%) | 15 | 6.7% (95% CI: -6.4–  19.7%) | 0% (IQR: 0–10%) |
| South Asia | 21 | 15.5% (95% CI: 3.2–  27.8%) | 7.7% (IQR: 0–  28.6%) | 10 | 11.8% (95% CI:  10.4–13.2%) | 11.8% (IQR:  11.4–12.2%) | 13 | 18.8% (95% CI:  10.2–27.5%) | 16.7% (IQR:  14.6–22%) |
| Pacific | 21 | 19.7% (95% CI:  12.6–26.7%) | 18.2% (IQR:  16.7–22.2%) | 6 | 0% (95% CI: 0–0%) | 0% (IQR: 0–0%) | 9 | 41.1% (95% CI: -6–  88.2%) | 40% (IQR: 20–  61.6%) |
