## Supplementary material for "Tracking two decades of global dengue virus lineage dynamics": S3 Table

### S3 Table. Summary of major and minor dengue lineages

| **Lineage** | **No. Sequences** | **Regions Found** | **Year First Detected** | **Region First Detected** |
| --- | --- | --- | --- | --- |
| ***Major Lineages*** |  |  |  |  |
| 1I_K | 4008 | North America, South America, Europe, SSA, East Asia, Southeast Asia, South Asia, Pacific | 2000 | Southeast Asia |
| 1III_A | 2825 | North America, South America, Europe, SSA, MENA, East Asia, Southeast Asia, South Asia, Pacific | 2004 | South Asia |
| 1IV_B | 650 | North America, South America, Europe, East Asia, Southeast Asia, South Asia, Pacific | 2000 | Southeast Asia, Pacific |
| 2II_A | 2061 | North America, Europe, SSA, MENA, East Asia, Southeast Asia, South Asia, Pacific | 2001 | Southeast Asia, South Asia |
| 2II_F | 4586 | North America, South America, Europe, MENA, East Asia, Southeast Asia, South Asia, Pacific | 2000 | Southeast Asia |
| 2III_C | 893 | North America, South America, Europe, SSA, East Asia, Southeast Asia, Pacific | 2000 | South America |
| 3I_A | 975 | North America, South America, Europe, East Asia, Southeast Asia, South Asia, Pacific | 2000 | Southeast Asia |
| 3III_A | 298 | South America, SSA, MENA, East Asia, Southeast Asia, South Asia, Pacific | 2003 | South Asia |
| 3III_B | 2137 | North America, South America, Europe, SSA, MENA, East Asia, Southeast Asia, South Asia, Pacific | 2002 | Europe, Southeast Asia |
| ***Minor Lineages*** |  |  |  |  |
| 1I_K.3 | 452 | South America, Europe, SSA, East Asia, Southeast Asia, South Asia, Pacific | 2010 | Southeast Asia |
| 1III_A.3 | 85 | North America, Europe, SSA, MENA, East Asia, Southeast Asia, South Asia | 2013 | Southeast Asia |
| 2II_A.1 | 1192 | North America, Europe, SSA, MENA, East Asia, Southeast Asia, South Asia, Pacific | 2001 | Southeast Asia, South Asia |
| 2II_F.1 | 4058 | North America, South America, Europe, MENA, East Asia, Southeast Asia, South Asia, Pacific | 2005 | Southeast Asia |
| 3III_B.3 | 1414 | North America, South America, Europe, SSA, MENA, East Asia, Southeast Asia, South Asia, Pacific | 2002 | Europe |
