## Supplementary figures and images for "Tracking two decades of global dengue virus lineage dynamics"

### S1 Fig

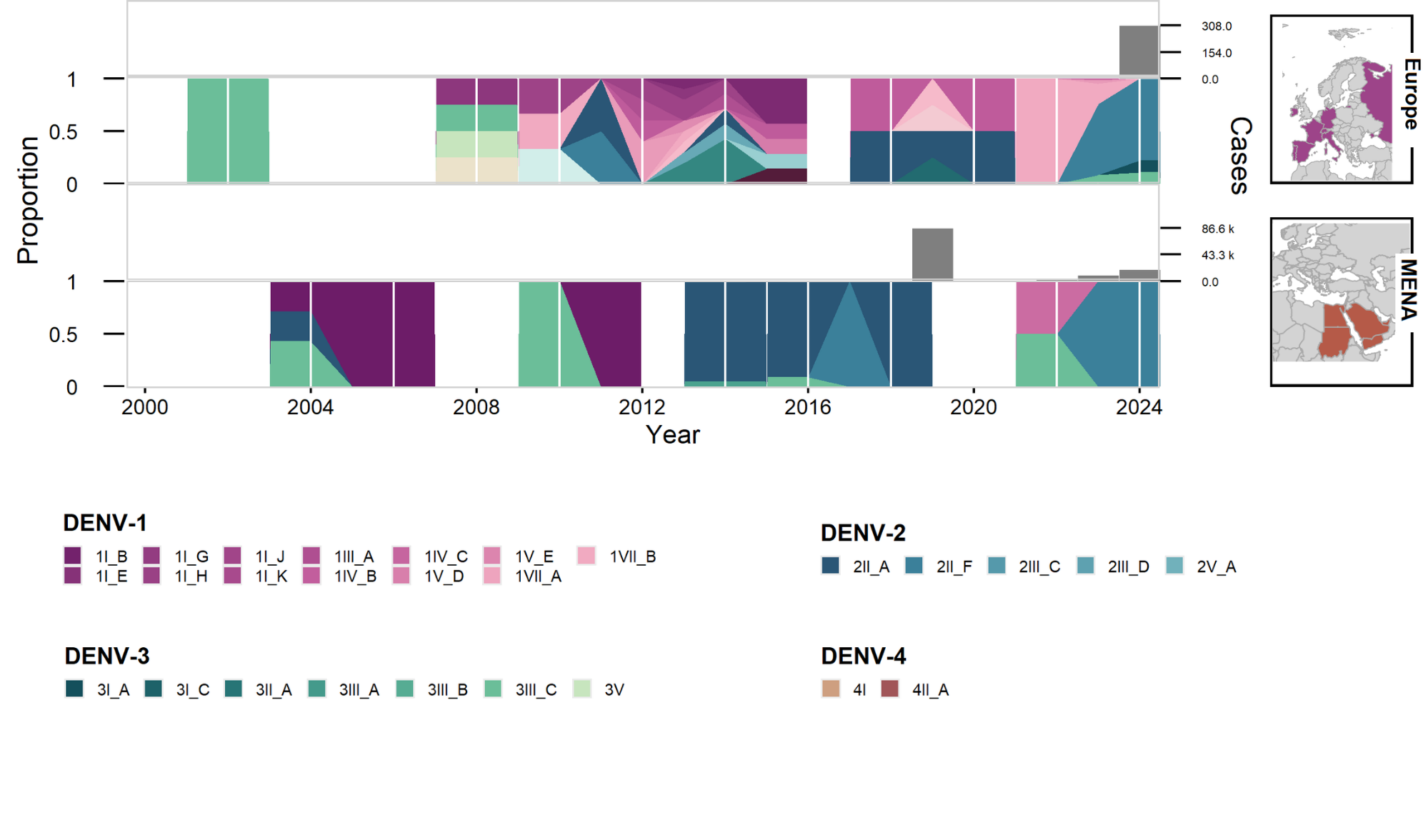

### S2 Fig

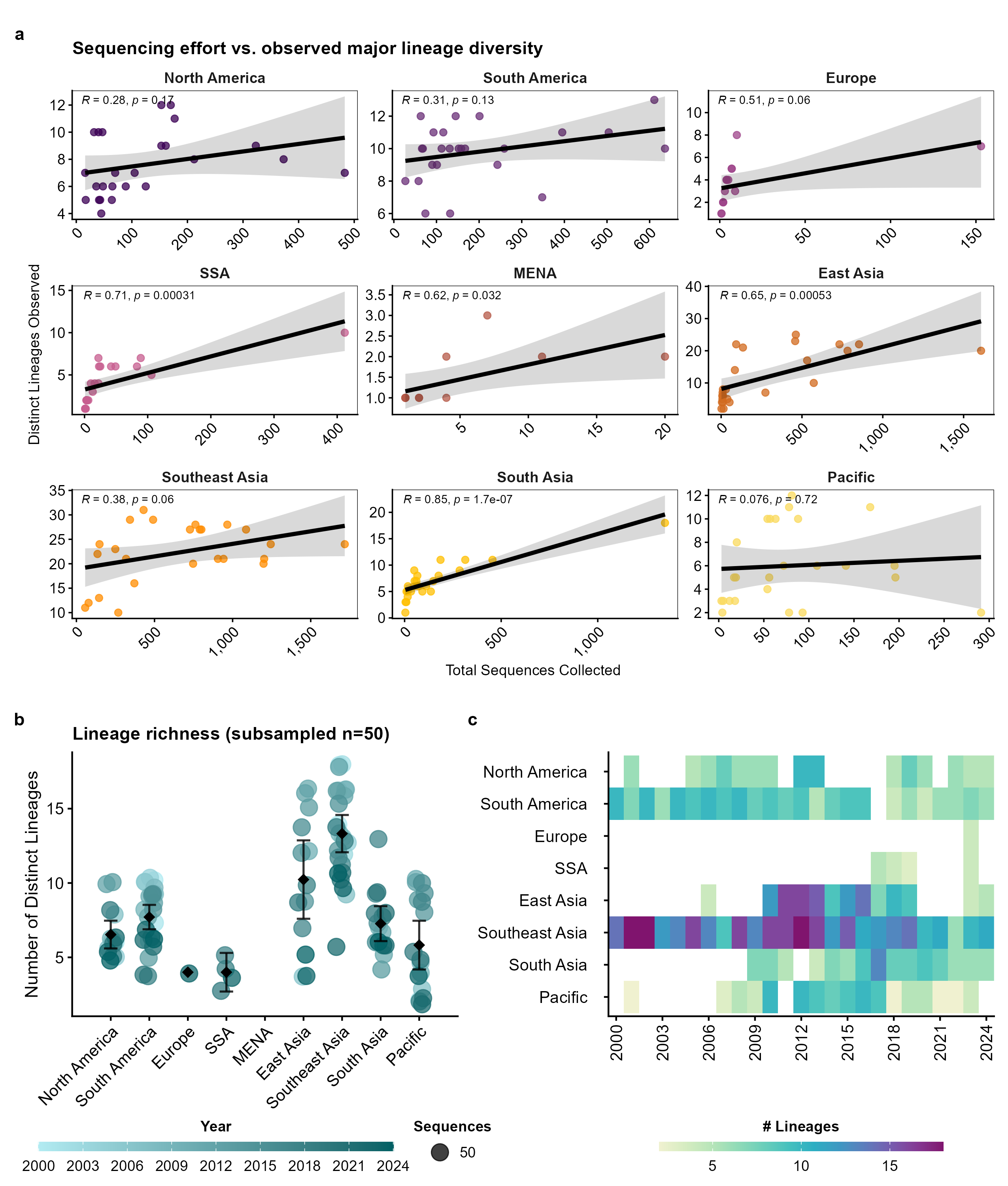

### S3 Fig

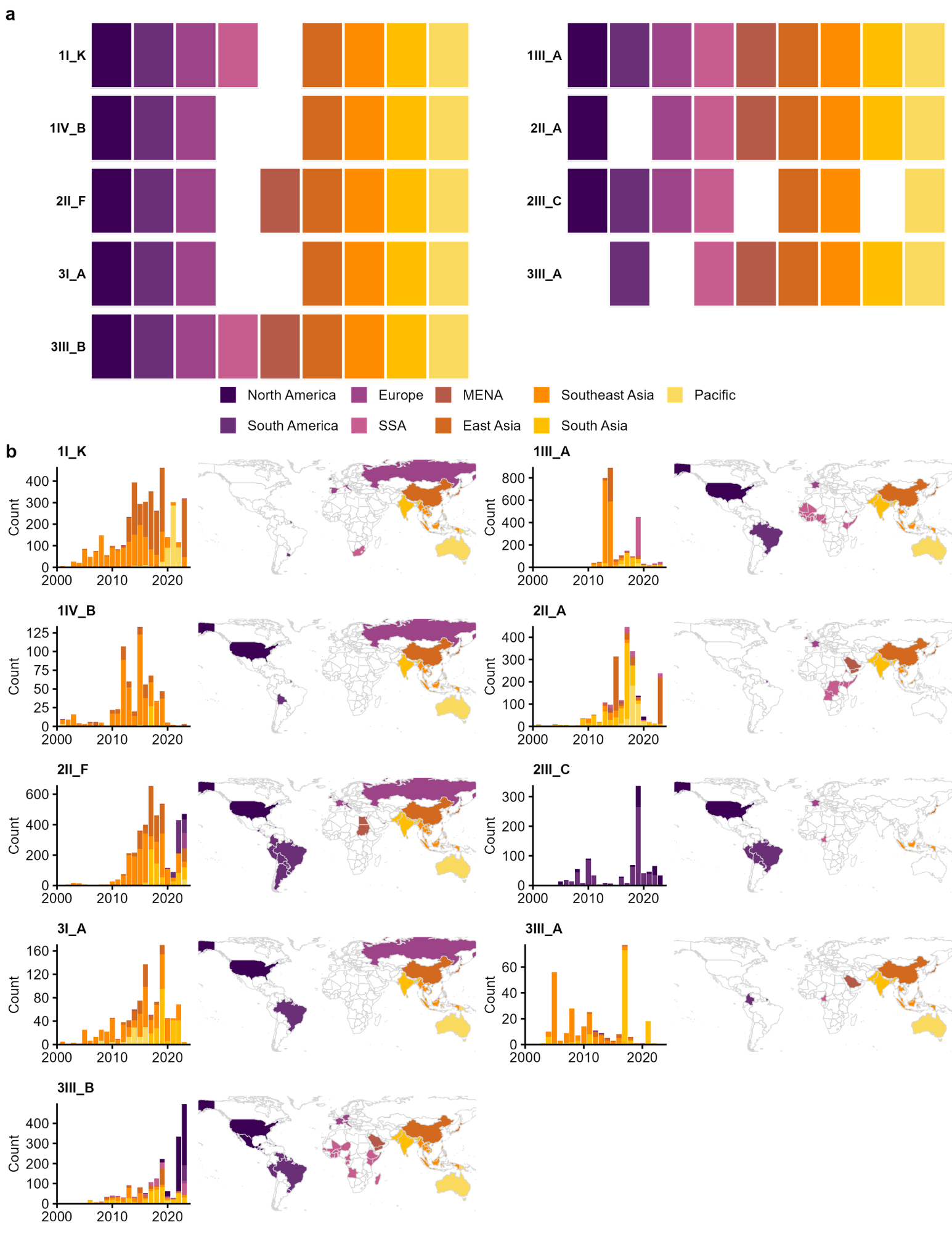
